# Fast, versatile, and quantitative annotation of complex images

**DOI:** 10.1101/391169

**Authors:** Kathleen Bates, Shen Jiang, Ruth Bates, Shivesh Chaudhary, Emily Jackson-Holmes, Melinda Jue, Erin McCaskey, Daniel Goldman, Hang Lu

## Abstract

We report a generic smartphone app for quantitative annotation of complex images. The app is simple enough to be used by children, and annotation tasks are distributed across app users, contributing to efficient annotation. We demonstrate its flexibility and speed by annotating >30,000 images, including features of rice root growth and structure, stem cell aggregate morphology, and complex worm (*C. elegans*) postures, for which we show that the speed of annotation is >130-fold faster than state-of-the-art techniques with similar accuracy.

## Introduction

The accelerating ease of collecting very large image data sets (terabytes to petabytes) has led to a shift in scientific bottlenecks from image collection to image analysis across many disciplines, including connectomics^1–3^, cell lineage tracing^4^, and ethology^5,6^. Although highly specialized computational pipelines are emerging to address this new bottleneck, these pipelines require significant effort to develop, are computationally expensive and not error-free, and may still rely on human image annotation. The widespread dependence on human image annotation or correction is likely to continue, and yet tools for image annotation, especially at large scales, often do not meet the needs of researchers.

Specifically, tools for quantitative annotation of images are hindered primarily by a trade-off between speed, accuracy, and versatility. Some tools require extensive tuning or parameter optimization for accurate annotation, or may not be well-suited for heterogeneous image quality. In addition, many tools limit the way users can define image features of interest, for example, via rectangles, polygons, or circles. Annotation speed is limited by the complexity of annotation software, and, ultimately, how quickly annotators can mark phenotypes accurately^7^. Equally critical for efficient annotation of large datasets is ease in distributing annotation tasks, as well as broadness in settings or locations where users can annotate. To serve the greatest number of researchers effectively, tools for large scale image annotation should be generalizable, fast, and accurate.

Here we report a highly versatile, fast, and quantitative method for image annotation. Features of interest of an arbitrary image can be annotated simply from user’s finger- or stylus-tracings (**Supplemental Movie 1**). We demonstrate the use of a simple and intuitive smartphone- and tablet-based app to annotate complex body postures in *Caenorhabditis elegans,* morphology of stem cell aggregates, and root growth of *Oryza sativa* (rice) and *Zea mays* (corn). We crowd-sourced annotations of over 16,000 nematode images, 500 stem cell aggregate images, and 900 root images, with a total of over 30,000 user annotations (**Fig. 1a-e**). Briefly, the app loads images from an online database to the user’s Android device, on which users draw their best annotation. The user then uploads the annotated image (as well as pixel vectors) and is immediately presented with another image from the image set. Our worm tracing example app, ‘Wurm Paint’, can be found for free on the <u>Google Play Store</u>, and the source code as well as setup instructions for creating new versions of the app can be found at https://github.com/jiangshen/WurmPaint.

**Figure 1.**
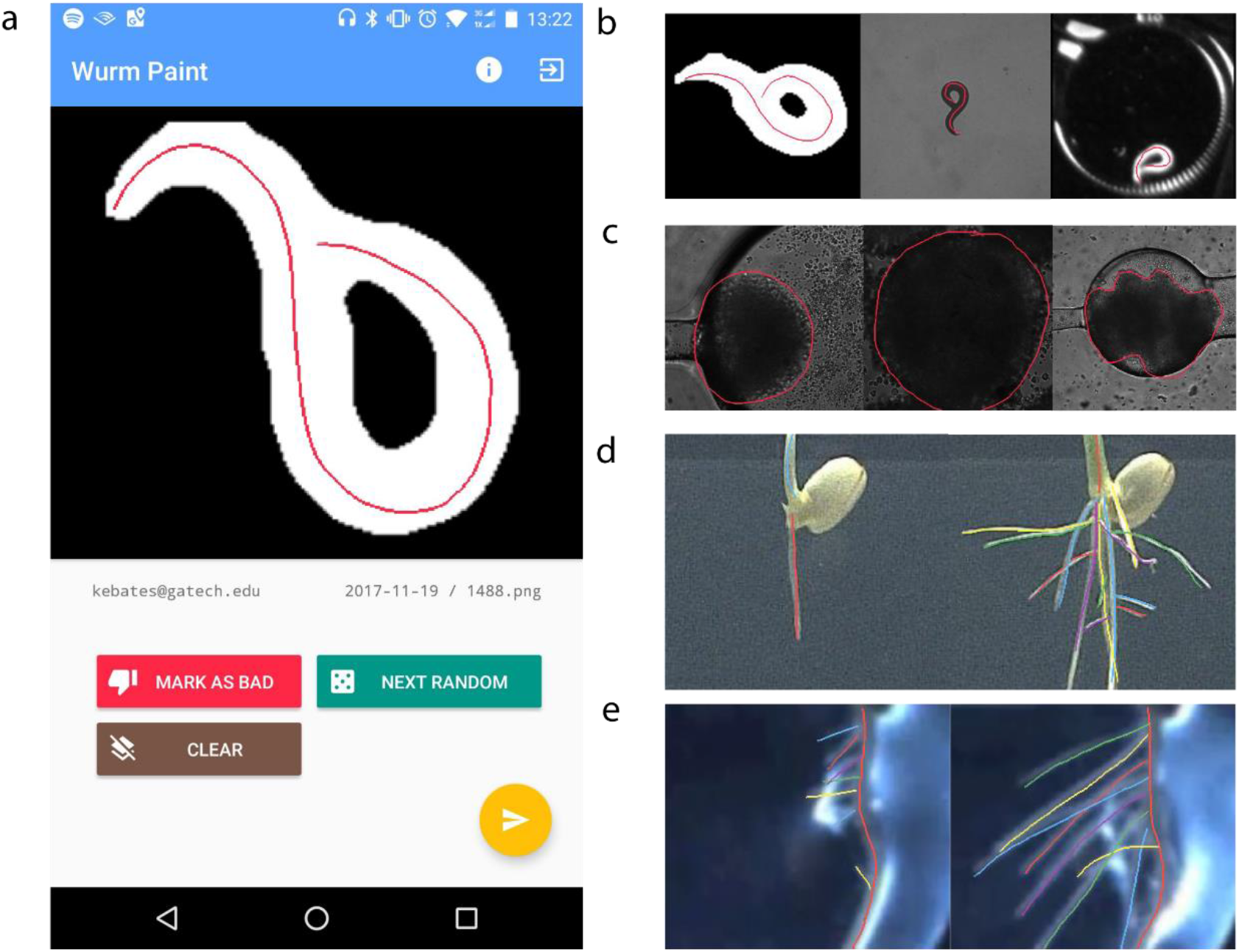
Versatile smartphone annotation of images. (**a**) Screen capture of Android interface of worm tracing app, ‘Wurm Paint’. See **Supplemental Movie 1** for a video of the app in use. (**b**) User annotations of worm posture in binary image, grayscale brightfield image, and grayscale darkfield image. (**c**) User annotations of stem cell aggregate morphology using app with same source code as Wurm Paint. (**d**) User annotations of rice root structure. Left-hand images temporally precede right-hand images. App presents images in temporal order, as temporal context helps users to decipher additional information when root structure is partially occluded at later timepoints. (**e**) User annotations of corn root structure. Left-hand images temporally precede right-hand images. Even in poor, changing, or inhomogeneous imaging conditions, human annotators make reasonable predictions.

## Results and Discussion

Our app is indiscriminate to the nature of images or annotations. Worm images on our database were derived from brightfield and darkfield microscope configurations, solid and liquid imaging environments, and included both processed, binarized images as well as unprocessed frames from raw videos (**Fig 1b**). Stem cell aggregate images on our database were derived from phase images of both live and fixed aggregates grown in tissue culture plates as well as aggregates grown in microfluidic devices (**Fig 1c**). For both nematode and stem cell aggregate applications, users are presented with randomized images from the full dataset and draw a single contour. This generic annotation scheme could also be used to trace individual cells, or features of developing embryos (such as *Drosophila melanogaster, Xenopus*, or zebrafish), to name a few. To allow users to annotate video frames in a pre-defined order (e.g. when temporal context is critical to annotation) and in cases where an image contains multiple features of interest, we created a second version of the app that presents uploaded images in order and allows users to draw as many contours as needed. We used this app version to annotate rice and corn root systems (**Fig 1d-e**). We expect that these two versions of the app could serve many other image annotation problems equally well with little to no changes of the source code.

The app is extremely easy for annotators to use. By using smartphones as the basis for our image annotation system, users need only draw with a finger or stylus, as compared to the greater difficulty of drawing with a computer mouse. The interface itself is simple and intuitive compared to popular image annotation and analysis tools. We had 7-12 year-olds use the worm tracing app, and found that it was simple enough for them to use without help after a brief explanation (**Fig 2a**). Although the quality of children’s annotations was far more variable than annotations by adults, many of the children’s annotations were of indistinguishable quality compared to those of adults and annotations inconsistent with other user’s annotations were easy to identify.

**Figure 2.**
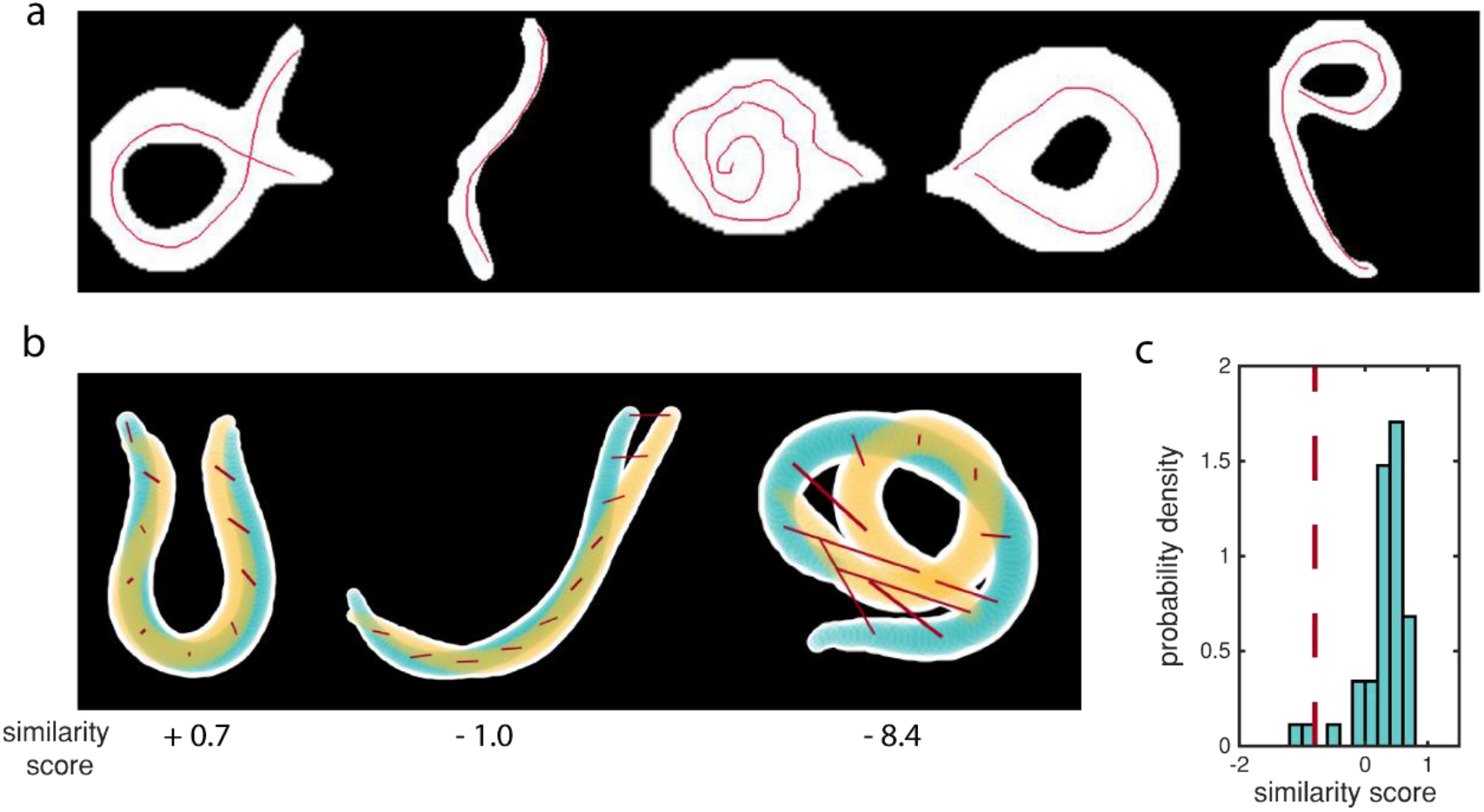
Intuitive and accurate annotation of images. (**a**) Annotations by 7-12 year-olds. Although annotation quality is variable, the app is simple enough that children could use it without any aid. (**b**) Sketch of similarity score calculations. For each panel, two worm contours (white overlaid with yellow or blue) are reconstructed based on ground truth or annotated midlines. 100 points along the midline are matched, and the Euclidean distance between each pair is computed. Here we show this at 10 points along the backbone (red lines). We then normalize the sum of these distances so that identical midlines result in similarity scores of one and scores of zero indicate that midlines were on average within three-quarters of the worm width apart. Yellow and blue highlight the center three-quarters of the worm’s width. (**c**) Probability density of similarity scores for unambiguous posture solutions compared to averaged user annotations of the same unambiguous postures. Most similarity scores lie above zero, indicating that users can accurately annotate unambiguous images. The dashed red line indicates the threshold we use to calculate consensus contours. Almost all average user annotations are above this threshold. N = 44.

Next, we sought to demonstrate that our app enabled fast annotation. For two of our applications, we quantified the time between image uploads of single users as a conservative estimate of time per annotation. For worm tracing, which always required a single user-drawn contour, the average annotation time was 7 ± 0 s/image (95% CI), and for root tracing, which often required multiple contours per image, the average annotation time was 14 ± 1 s/image (95% CI). To benchmark user annotation speed in our app, we annotated worm images using ImageJ^8^, which routinely required more time. In addition to the importance of individual users’ speed, overall speed is dependent on how many users can annotate in parallel. Smartphone-based annotation not only allows us to easily distribute image annotation tasks as narrowly (a single expert) or broadly (general public) as desired, it also expands geospatial locations and settings where users can annotate^9^.

We then assessed the ability of users to trace known shapes accurately. We did this by comparing averaged hand-drawn worm postures to computationally generated ground truth postures. For worms with unambiguous postures, we matched points along the averaged hand-drawn worm midlines with points along the corresponding ground truth midline and summed the Euclidean norm of all point pairs. To determine an overall similarity between any two worm midlines, we reasoned that an acceptably similar midline should lie within the center three-quarters of the worm’s total width at any given point. We therefore normalized similarity scores so that a score of one indicated identical midlines, any positive score indicated that the midlines were on average less than three-quarters of the worm width apart, and negative similarity scores indicated that midlines were further than three-quarters of the worm width apart (**Fig 2b**). Most averaged annotations of unambiguous postures had similarity scores above zero when compared to their corresponding ground truth midline, including data collected from non-expert annotators (**Fig 2c**). We concluded that the annotation accuracy was sufficiently high for tracing worms.

To further demonstrate a practical application of our app, we focused on using annotations of ambiguous *C. elegans* postures to reconstruct the dynamics of worm behavior. Ambiguous postures result from segmentation errors, or more frequently, the worm partially occluding itself, for example during stereotyped Ω- or δ-turns. A major advantage of using human annotators is the ability to quickly generate varied predictions for images that humans and algorithms alike struggle to find a ground truth for. *C. elegans* postures are often simplistic and sinusoidal, but ^~^7% of the worms’ behavior results in postures that are impossible to segment using current tools. One approach relies on computationally expensive optimization to attempt a quantitative posture description^10,11^. Although accurate in most instances, this state-of-the-art strategy for predicting ambiguous nematode posture requires on average 931.7s (n = 66) per video frame (software configuration in supplemental). Based on our average worm annotation time, users can make predictions about 130-fold faster than this computational strategy. User predictions for individual ambiguous images varied, but could typically be grouped into several distinct shapes, indicating that there were often only a few reasonable predictions for each ambiguous posture (**Fig 3a**). To characterize this variability quantitatively, we calculated pairwise similarity scores comparing different annotations of the same image for more than 500 source images and found that similarity scores peaked between zero and one, and had a left-skewed distribution with a significant tail (**Fig 3b**). This is consistent with our observation that although there is significant variability in user annotations, users are frequently in agreement with one another, suggesting the utility of a consensus-based approach in identifying a best solution. The ease and speed of generating viable predictions based on human intelligence with the app gives it particular advantage in analyzing images where a single ‘correct’ solution is non-existent and several solutions have high likelihood.

**Figure 3.**
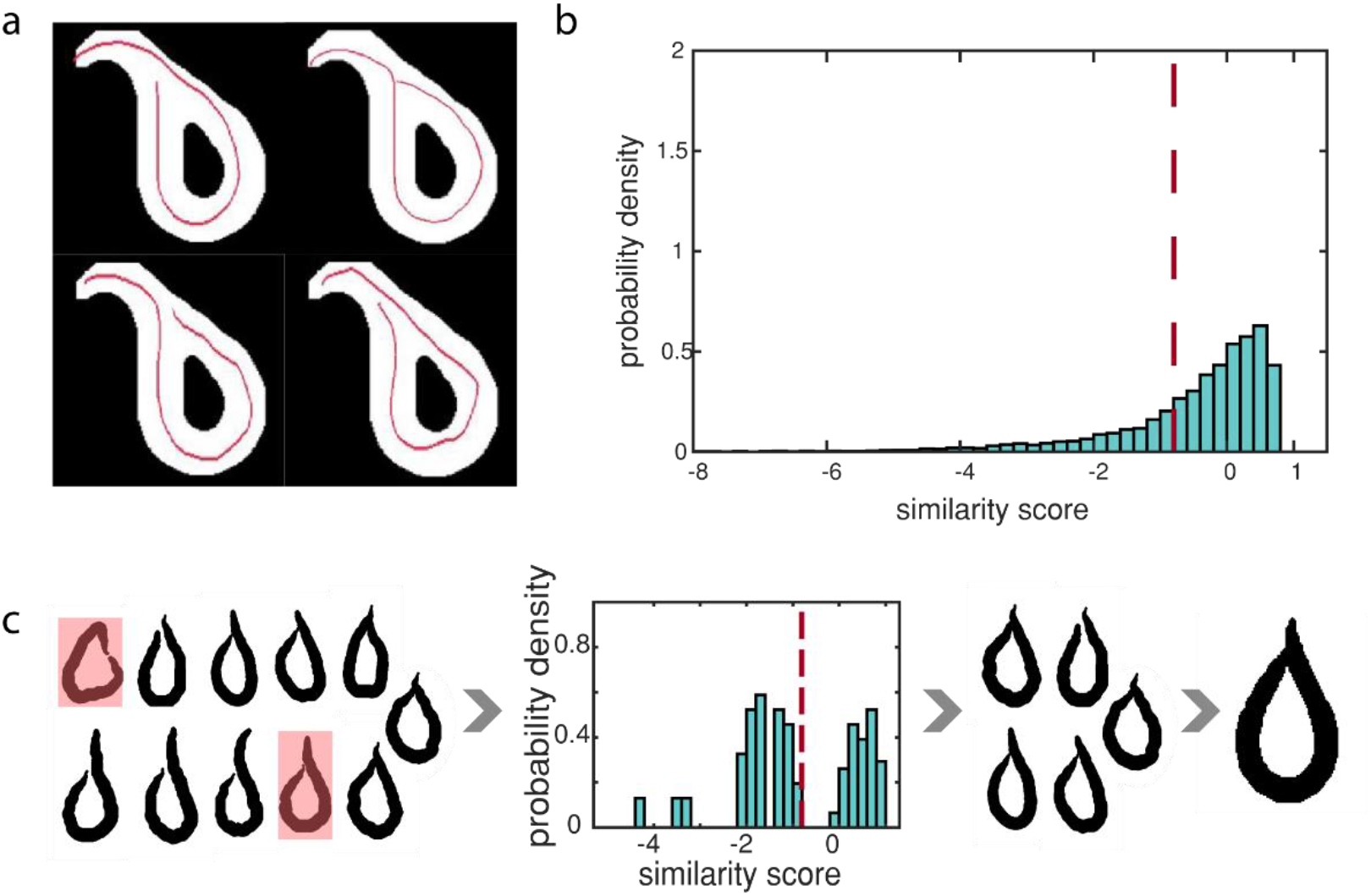
Generating accurate consensus predictions for ambiguous images. (**a**) Example set of ambiguous images. While some users draw the midline as continuous to the left side of the worm body, others draw the midline as continuous to the right side of the worm body. (**b**) Probability density of similarity scores for comparison between different user annotations of the same ambiguous posture. The skew of the probability density is consistent with higher variability in user annotations of ambiguous postures compared to unambiguous postures, while the mode location between zero and one is consistent with many users in agreement with one another. The dotted red line indicates the threshold we use to calculate consensus contours. Although this threshold is less than zero (i.e. midlines are greater than three-quarters width distant), annotations above this threshold are generally consistent with one another. The threshold was determined by modeling probability density as a mixture of two Gaussians (see **Methods**). N =26,098. (**c**) Illustration of consensus generation scheme. First, we identify and remove outlier annotations and annotations outside of a range of eigenmode amplitudes. Next, we compare each annotation to every other annotation of the same posture using similarity scores. We group annotations based on which pairs have similarity scores above the dotted red threshold. Then we find the group containing the most annotations and use the cluster centroid to produce a consensus contour.

To resolve the ambiguities in our postural data set, we used annotations to create a consensus prediction for ambiguous images (**Fig 3c**). For each source image, we first eliminated annotations that were outliers or that created shapes outside of *C. elegans* postural space, then used pairwise similarity scores to identify groups of similar annotations. We then chose the group containing the most individual annotations, and averaged annotations in this group to come to a consensus contour. Finally, we compared these disambiguated annotations to predictions generated by the state-of-the-art computational method and found that the mode of the similarity score distribution was −1, indicating that although consensus contours had somewhat reduced accuracy, they overall agreed well with computational predictions (**Supplemental Figure 2b**). Further, for frames where initial segmentation failed, users could correctly annotate grayscale source images, while computational predictions were erroneous.

*C. elegans* is a powerful model organism with a large suite of tools for genetic manipulation^12,13^. These tools, along with a fully mapped nervous system^14^, have enabled researchers to identify molecular mechanisms and individual genes associated with behavioral phenotypes^15–17^. However, quantitative analysis of some of the most complex behaviors, large-angle turns that commonly include ambiguous postures, remains difficult, and gaps in quantifiable behavior prevent dynamic posture analysis altogether. Using our consensus worm contours, we recreated the postural repertoire and behavioral dynamics of *C. elegans*. First, we sought to answer how significantly complex worm postures affect the overall shape space of *C. elegans*. To answer this, we calculated the first four principle components of *C. elegans’* shape space^18^ (‘eigenworms’) using either unambiguous results alone or both unambiguous results and consensus contours (**Fig 4a, Supplemental Figure 2c**). Consistent with prior reports, we found that the first four principle components were very similar with or without ambiguous postures^11^. Interestingly, the fractional variance of the worm’s posture space captured by these eigenworms is greater when ambiguous postures are included (**Fig 4b**). Lastly, we recreated complete timeseries of the first four eigenworm amplitudes for individual worms using the consensus contours (**Fig 4c**). These traces fill in the gaps left by ambiguous shapes and outperform the computational prediction in some cases where the worm is tightly coiled **(Supplemental Movie 2)**. In addition to adding to our knowledge of *C. elegans* behavioral dynamics purely through image annotation, this app can help improve existing posture prediction algorithms by using these results.

**Figure 4.**
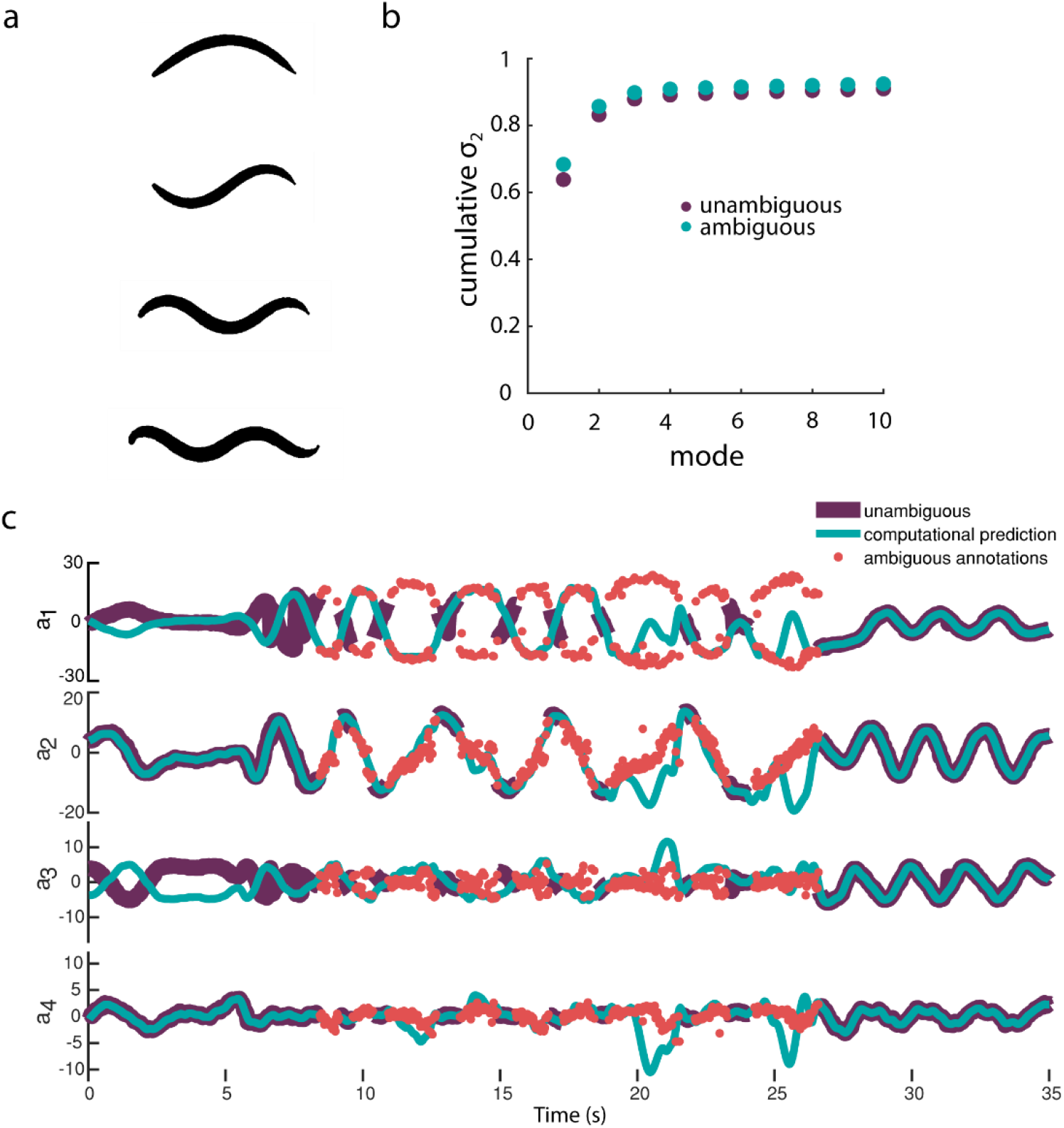
Reconstruction of continuous behavior dynamics from annotations. (**a**) Representation of first four principle components of worm posture (‘eigenworms’) including both unambiguous and ambiguous postures from four annotated videos in our dataset (37,784 frames). Our eigenworms are different from many other reported eigenworms, but this is a result of the extended physical environments represented in our dataset. As reported in other works, our eigenworms both with and without ambiguous postures were very similar (see **Supplemental Figure 2c**). (**b**) Cumulative variance captured by each additional eigenworm (‘mode’) for both unambiguous images only and both unambiguous and ambiguous images together. As in other work, we note that most of the variability in *C. elegans’* posture is captured by very few eigenworms, and after the first four eigenworms the additional variance captured is incremental for both ambiguous and unambiguous cases. When both unambiguous and ambiguous images are included, the variance captured by the first four eigenworms is greater than the variance captured using only unambiguous images. (**c**) Traces of amplitude of first four eigenworms in time for an individual worm. Dark purple lines are amplitudes calculated for unambiguous postures via image processing. Note the extensive gaps between purple lines that represent ambiguous postures that are usually associated with reorientation and turning of the worm. Blue lines are computational predictions for the full video, including ambiguous postures. Red dots represent consensus contours for individual frames found using app user annotations. Typically, red dots follow similar contours to blue lines. For the amplitudes of the first eigenworm in particular, the red dots follow two opposing sinusoidal contours simultaneously, one contour representing the opposite head orientation of the worm compared to the other contour. As is apparent in **Supplemental Movie 2**, there are some cases where user annotations more accurately represent the worm’s motion compared to the computational prediction.

Our app-based annotation scheme allows researchers from any field to quickly and easily annotate complex images in quantitative ways. Here, we demonstrated its flexibility and speed in annotating rice root growth and structure, stem cell aggregate morphology, and complex worm postures, where we showed that the app is ^~^ 130-fold faster than state-of-the-art posture optimization techniques. We expect that the app will be useful as an alternative to creating complex and bespoke computational image processing pipelines, as a way to complement and augment existing computational pipelines, and as a simple way to generate consensus ground truths towards improving machine learning algorithms for image processing.

## Materials and Methods

### *C. elegans* maintenance

Worms were cultured at 20C in a dark incubator on standard nematode growth medium (NGM) petri dishes seeded with OP50 bacteria^19^. All experiments were performed on day 1 adults. Animals were synchronized via 2 hr lay-offs; 4 days before experiments, ^~^ 20 adult worms were picked onto seeded plates and allowed to lay eggs for 2 hrs before being picked back off. Strains used in this work include N2; QL142: spe-27(it110); and AQ2334: lite-1(ce314); ljIs123[pmec-4 ChR2; punc-122 RFP].

### *C. elegans* behavior experiments

For on-plate experiments, in addition to age synchronization as described above, worms were cultured on NGM agar plates seeded with OP50 E. Coli containing either 100 microM ATR (Sigma-Aldrich) (experimental) or no ATR (control) until day 1 of adulthood. Experiments were performed on a previously developed microscopy system^20^ that tracks and records free-ranging behavior of individuals while optogenetically stimulating animals. For ‘swimming’ experiments, microfluidic devices were made as described^21^. Silicone tubing and metal luer connectors were rinsed with dH20 and autoclave sterilized. Devices were flushed and degassed with filtered and autoclaved M9 containing 0.01% Triton-X 100. After degassing, synchronized day 1 adult worms were washed off of agar culture plates and suspended in M9 containing 0.01% Triton-X 100. The worm suspension was loaded into device as described^21^. After loading, bulk flow within the device was prevented via medical clips closed on the silicone tubing.

### Android App and **database**

The Android application was written in Java and it utilizes Firebase as the database backend. The database structure can be found in **Supplemental Figure 1**. The app also makes use of Google Play Services to facilitate login via Google account and for gaming purposes. The app conforms to material design and focuses on clean user interfaces for better usability and smoother drawing experience. Code for the app can be found at https://github.com/jiangshen/WurmPaint, along with instructions on how to setup the app from the source code. The source code can be used under an MIT license.

To familiarize non-expert users with typical worm movement and shapes, we assembled a brief tutorial https://sites.google.com/view/wurm/tutorial. As general guidelines, we asked users to draw a continuous contour along the midline of the worm, starting at one end of the worm to the other end, so that the contour did not contain sharp corners, rather smooth bends along the length of the worm.

### Worm tracking

We built upon an existing worm tracker^11^ for our initial image analysis and to identify frames where worms were partially self-occluded (i.e. ambiguous). A subset of these frames was uploaded to our database for annotation. Using the generative algorithm included in the existing worm tracker to predict worm posture for occluded shapes, we optimized parameters for our data set and found predicted worm postures for several full videos from which we had drawn ambiguous postures for our database. The worm tracker uses MATLAB software (we used MATLAB version 2017a). Further software configuration and parameters we used can be found on our lab GitHub. To evaluate the time required to process individual frames using this worm tracker, we used MATLAB to measure how long the point-swarm (PS) optimization (generation of alternative posture predictions) required for each ambiguous frame. This step took an average of 776.2 ± 5.2 s/ ambiguous frame (95% CI, n = 66) with parallel processing (a local pool consisting of 4 cores). After this generative step, a progressive optimizing interpolation (POI) step evaluates the alternative posture predictions to determine which makes sense in the context of the worm postures in the surrounding frames. For this step, we timed the total time until a solution was generated. For a movie with 444 ambiguous frames, this step required 155.5 s/ ambiguous frame (equivalent to the time required for > 20 human annotations). Combined, the PS and POI tracking steps required on average 931.7 s/ ambiguous frame, or the equivalent of 133 human annotations. The computers used were Dell Precision Tower model 5810 with 32 GB RAM and Intel Xeon CPU.

### User annotation speed

We collect timestamps when users upload images and drawing vectors with a resolution of 1s, based on the user’s device’s time. To determine a conservative average user annotation speed, we first grouped all annotations by user, then computed the time between each upload for that user. We then pooled all these inter-upload times. Because inter-upload times could range from a few seconds to days depending on the user’s usage frequency, we imposed an upper threshold of 30s for worm image annotations and an upper threshold of 90s for root image annotations to determine the average user annotation speed.

### Post-processing of annotated worm images

Although the current version of the app allows us to upload the coordinate trajectories of user annotations, the initial version that much of the data presented here originates from only allowed us to upload the annotation superimposed on the source image. Thus, to extract annotations and reconstruct trajectories from uploaded images, some post-processing of annotated images was required. Briefly, to identify annotations, we found non-grey pixels in each image. We then binarized the annotation alone and skeletonized the image, followed by removal of branch points if branch points existed. We then checked the curvature of each line segment to ensure it fell in a reasonable range - if it did not, we broke the segment at its point of maximum curvature. Using the resulting line segments, we attempted to reconnect them to each other using both the proximity of segment endpoints and local segment slope. Once segments had been reconnected, the worm’s midline was reconstructed using the projections onto the first five eigenvectors as described previously^18^. Average speed of this post-processing was 0.0597 s/ frame (n = 1000). This process is illustrated in **Supplemental Figure 2a**, and code for these steps is available in our GitHub repository. However, we emphasize that other app users need not perform any post-processing of images. Instead, coordinate trajectories can be accessed by parsing JSON files that are downloadable from our Firebase database.

### Similarity score calculation

As described in the text, in order to compare two paths, we matched 100 points between two worm midlines and computed the Euclidean distance between each pair, followed by summing all of these distances and normalizing the distance by 75% of the width of that particular worm at each of the 100 matched points. Mathematically, 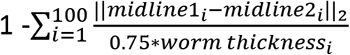.

### Consensus generation

To construct consensus midlines from user annotations, we first noted that even for pairs of reconstructed midlines that were below a zero similarity score, users were making essentially the same annotation. To identify a threshold similarity score below which we could consider two annotations to be from distinct groups, we modeled the distribution of similarity scores from user-user comparisons (**Fig 2d**) as a mixture of gaussians. The primary mode was centered at −0.068 and the secondary mode was centered at −3.260 **(Supplemental Figure 2b).** To ensure that most generally similar annotations were grouped together, we computed a threshold two standard deviations below the primary mode, a similarity score value of −0.809. We found that several other methods of identifying this similarity score threshold identified thresholds that ranged from slightly positive to slightly negative. These methods included the Otsu thresholding method on user-user similarity scores and searching for the lowest threshold of the user-user similarity scores for which the Wilcoxon rank-sum test failed to reject the null hypothesis that the user-user similarity scores and user-ground truth similarity scores were drawn from the same distribution at the 5% significance level.

Having identified a reasonable threshold, we generated consensus contours. During this process, we used the projections of worm backbones into the space of the first five eigenvectors. First, we identified and removed annotations whose eigenvector projections were outside of the range of *C. elegans* posture space. Then, for each source image, we first identified all annotations of the source image and removed any remaining outlier annotations of that image, where an outlier is a value more than three scaled median absolute deviations away from the median. Next, we computed similarity scores for all pairs of annotations and used our previously identified threshold to identify pairs of images that were very similar to one another. Then, we further grouped these pairs into larger groups of similar annotations and identified the group of similar annotations with the largest number of members. For example, if image pairs (1, 2), (2, 3) and (5,6) all have similarity scores above our threshold, we take the union of all pairs that contain images 1, 2 and 3 and, separately, the union of all pairs that contain images 5 and 6. If more images belong to the first union set than the second, we use the first set to calculate a consensus contour by finding the centroid of this group of contours in the five-dimensional space of posture projections.

### Code Availability

Source code for Wurm Paint and Root Trace (root tracing app) are provided at https://github.com/jiangshen/WurmPaint. along with instructions for independent researchers to create their own versions of the app using novel data. The source code is available under an MIT license. Analysis code is available at https://github.gatech.edu/Lu-Fluidics-Lab/WurmPaintProcessing (with dependency on https://github.gatech.edu/Lu-Fluidics-Lab/eigenwormTracker).

### Data Availability

The datasets generated during and/or analysed during the current study are available from the corresponding author on reasonable request.

## Author Contributions

K.B. and H.L. conceived the project and wrote the manuscript, S.J. developed the app with input from K.B. and H.L., K.B. developed supporting code and analyzed data, D.G. and E.M. provided *O.sativa* data, E. J-H. provided stem cell aggregate data, K.B., H.L., R.B., M.J., E.J-H. and S.C. annotated data.

---

**Supplemental Movie 1** | Demonstration of example app usage. Users simply trace features of interest with a finger or stylus and upload them to the cloud. Users can also report images that don’t match the anticipated pattern (in this case, images that don’t look like worms). Movie is in real time.

**Supplemental Movie 2** | Representative example reconstruction of worm behavior. Segmented images from raw data are on left, with purely algorithmic reconstruction in center, and consensus annotation-based reconstruction on right (only for complex or ambiguous frames). Although algorithmic reconstruction is visually more accurate in many frames, annotation-based reconstruction produces accurate results even in frames that are inaccurately segmented or in cases where the algorithm performs poorly. Movie is at half speed from original 30 fps.

## Supplemental Figures

**Supplemental Figure 1.**
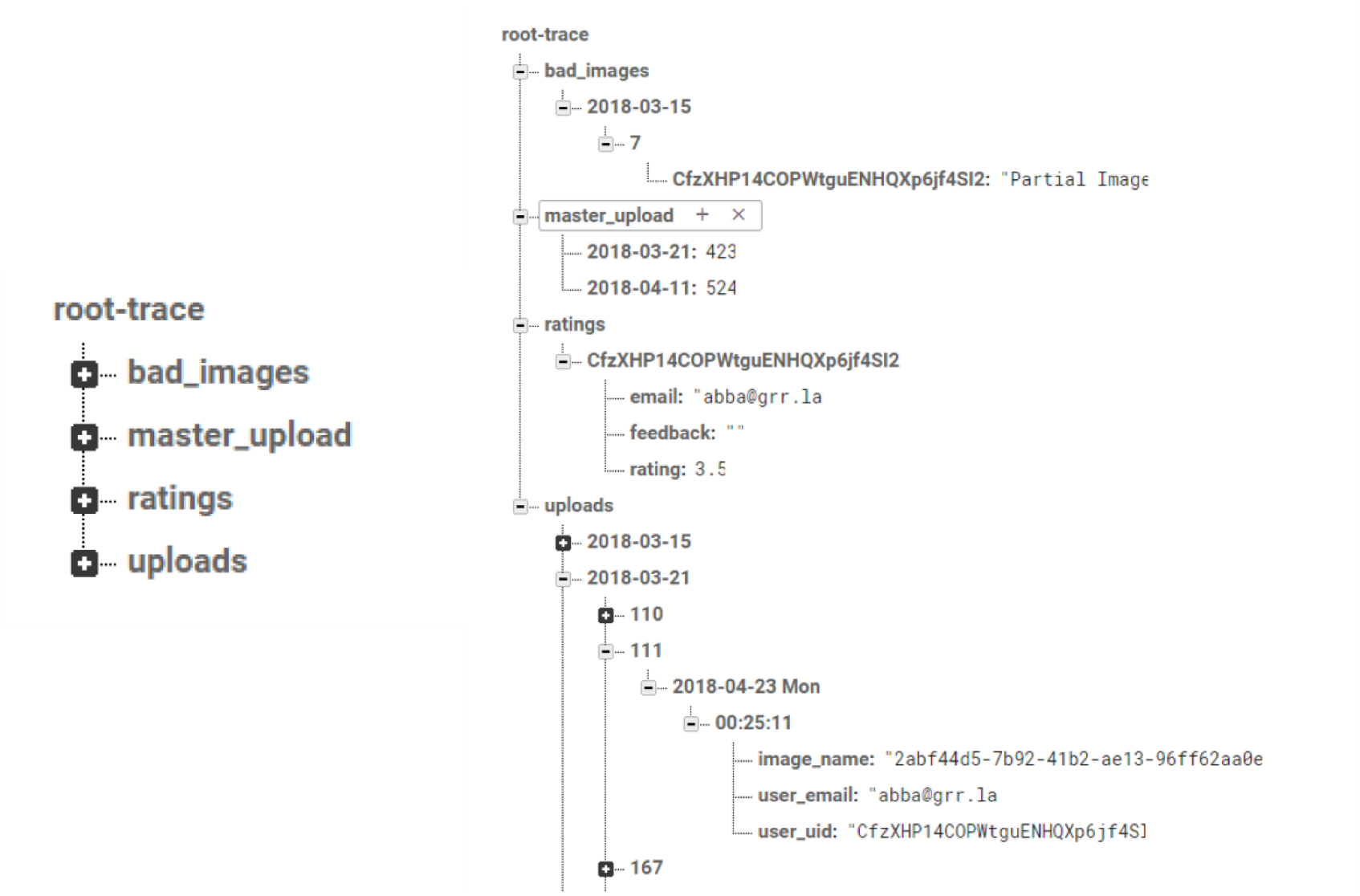
App database structure. Top-level (left) and expanded (right) structure of the root tracing app database. All apps have similarly structured databases. ‘Bad_images’ contains mapping to user-reported images. In Wurm Paint, we used this feature to identify images where image segmentation had failed, where the image feature was too small for users to draw, or images that didn’t clearly contain a worm. ‘Master_upload’ defines which source image sets are live on the app, as well as the number of images in each source set. User feedback is stored in the ‘ratings’ structure. Finally, ‘uploads’ maps user annotations (with user id, image name, and date and time of annotation) to the source image. In newer app versions available on our Github, we also save line trajectories at the bottom of the ‘uploads’ structure. To initialize the app, only the ‘master_upload’ structure is needed.

**Supplemental Figure 2.**
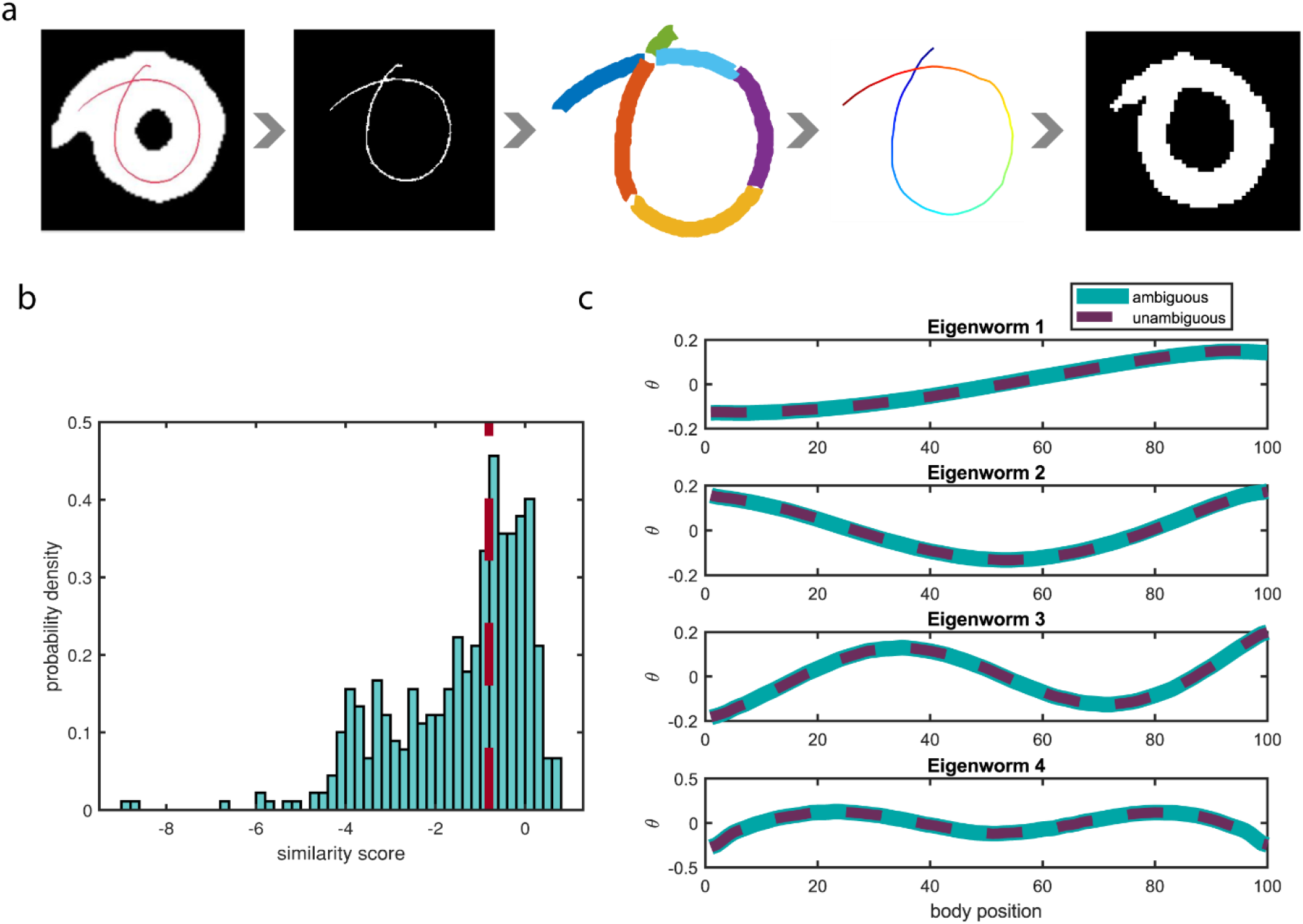
Worm annotation characterization (**a**) Reconstruction pipeline for worm midlines. Worm backbone annotations used in the main text were collected as images superimposed on the source image, so midlines must be reconstructed. Newer versions of the app save the drawn trajectory directly, so reconstruction is unnecessary. We found non-gray pixels (annotations) within the annotated images and binarized the annotation, followed by breaking the annotation at points of intersection or extreme curvature. Then we used local curvature and distance metrics to predict which line segments were connected and reconstructed the midline and image. (**b**) Probability density of similarity scores for ambiguous posture predictions compared to ambiguous posture consensus annotations. Red dashed line is threshold used for consensus generation. The broader similarity score distribution compared to unambiguous annotation to unambiguous ground truth comparison is caused partially by user variability and lower user accuracy and partially by the predictive nature of the state-of-the-art algorithm that sometimes leads to incorrect solutions. N = 449 (**c**) First four eigenvectors (‘eigenworms’) of the *C. elegans* posture space computed from four annotated videos (>37,000 frames). Computing eigenworms from only unambiguous postures or both ambiguous and unambiguous postures resulted in little difference, as reported in other work. Compared to eigenworms reported in other work, ours are similar, but with different eigenworms capturing a greater fraction of the total postural variability.

